# Multi-trait Multi-environment Genomic Prediction Strategies for *Miscanthus sacchariflorus* Populations

**DOI:** 10.64898/2026.03.18.712730

**Authors:** Shatabdi Proma, Garcia-Abadillo Julian, Vitor Sagae, Erik Sacks, Andrew D.B. Leakey, Hua Zhao, Bimal Kumar Ghimire, Alexander E. Lipka, Joyce N. Njuguna, Chang Yeon Yu, Eun Soo Seong, Ji Hye Yoo, Hironori Nagano, Kossonou G. Anzoua, Toshihiko Yamada, Pavel Chebukin, Xiaoli Jin, Lindsay V. Clark, Karen Koefoed Petersen, Junhua Peng, Andrey Sabitov, Elena Dzyubenko, Nicolay Dzyubenko, Katarzyna Glowacka, Moyses Nascimento, Ana Carolina Campana Nascimento, Maria S. Dwiyanti, Larisa Bagment, Ansari Shaik, Diego Jarquin

**Affiliations:** Agronomy Department, University of Florida, Gainesville, Florida, USA; Center for Advanced Bioenergy and Bioproducts Innovation, University of Illinois at Urbana Champaign, Urbana, IL 61801, USA; Department of Crop Sciences, University of Illinois at Urbana Champaign, Urbana, IL 61801, USA; Institute for Genomic Biology, University of Illinois at Urbana Champaign, Urbana, IL 61801, USA; Department of Plant Biology, University of Illinois at Urbana Champaign, Urbana, IL 61801, USA; Center for Digital Agriculture, University of Illinois at Urbana Champaign, Urbana, IL 61801, USA; College of Plant Science and Technology, Huazhong Agricultural University, Wuhan, China; Bio-Herb Research Center, Kangwon National University, Chuncheon 200-701, Korea; Division of Bioresource Sciences, Kangwon National University, Chuncheon, Korea; Field Science Center for Northern Biosphere, Hokkaido University, Sapporo, Japan; FSBSI “FSC of Agricultural Biotechnology of the Far East named after A.K. Chaiki”, Ussuriisk, Russian Federation; Key Laboratory of Crop Germplasm Research of Zhejiang Province, Agronomy Department, Zhejiang University, Hangzhou, China; Research Scientific Computing, Seattle Children’s Research Institute, Seattle, Washington, USA; Schroll Medical ApS, Arslev, Denmark; Spring Valley Agriscience Co. Ltd., Jinan, Shandong, China; Vavilov All-Russian Institute of Plant Genetic Resources, St. Petersburg, Russian Federation; Department of Biochemistry, University of Nebraska-Lincoln, Lincoln, Nebraska; Laboratory of Intelligence Computational and Statistical Learning (LICAE), Department of Statistics, Federal University of Viçosa, Viçosa, Brazil; Research Faculty of Agriculture, Hokkaido University, Japan; Cand. Sci (Biology), Leading Researcher, N.I. Vavilov All-Russian Institute of Plant Genetic Resources

**Keywords:** Genomic Prediction (GP), Genotype-by-Environment interaction, *Miscanthus sacchariflorus* (MSA), Multi-Trait Multi-Environment Prediction, Single Trait Multi-Environment Prediction

## Abstract

Genomic selection holds the potential to serve as a strategic tool to enhance the genetic gain of complex traits in *Miscanthus* breeding programs. The development of improved cultivars requires their assessment for various traits across diverse environments to ensure suitable overall performance. Hence, the multi-trait multi-environment (MTME) genomic prediction (GP) models offer an opportunity to improve selection accuracy. This study aims to evaluate the potential of five GP models: (1) three MTME models including genotype-by-trait-by-environment interaction (G×E×T) and (2) two single-trait multi-environment (STME) models (with and without G×E interaction). A *Miscanthus sacchariflorus* population comprising 336 genotypes evaluated in three environments and scored for four traits (biomass yield YDY, total culm number TCM, average internode length AIL, and culm node number CNN) was analyzed.

The predictive ability of the models was evaluated considering three cross-validation schemes resembling realistic scenarios (CV1: predicting new genotypes, CVP: predicting missing traits in a given environment, and CV2: predicting partially observed genotypes). On average, in all cross-validation schemes compared to the STME the predictive ability of the MTME models was 10% to 70% higher for TCM and AIL. On the other hand, for YDY and CNN, both STME models performed similarly or slightly better (between 5 to 64%) than the MTME models in most environments. While the MTME models were not successful for all traits when compared to their STME counterparts, MTME models improved the prediction of the performance of genotypes that were untested across environments or lacked trait information in a specific environment. Overall, our study suggests that MTME GP models can be implemented in *Miscanthus* breeding programs to improve the predictive ability of the complex traits, shorten breeding cycles, and accelerate selection decisions.

## Introduction

Plant biomass presents a significant source for domestic supply chains of sustainable bioenergy production (Saha et al., 2013). Plant species utilized for bioenergy applications have the potential to accumulate higher biomass with low inputs, thereby improving soil health and water quality (Clifton-Brown et al., 2008; Sacks et al., 2013). *Miscanthus*, a C4 perennial grass species, is a promising candidate for bioenergy production due to its high biomass yield with minimal inputs (Clifton-Brown et al., 2008; Clark et al., 2019). *Miscanthus* biomass can be used for paper production, direct combustion for heat and electricity generation, lignocellulosic ethanol production, animal bedding, and livestock feed (Clifton-Brown et al., 2001; Clifton-Brown and Lewandowski, 2002; Burner et al., 2017).

Currently, the commercial production of *Miscanthus* is limited to a sterile triploid clone, *Miscanthus* ×*giganteus* (M.×g), which is a cross between *Miscanthus sacchariflorus* (MSA) and *Miscanthus sinensis* (MSI) (Hodkinson and Renvoize, 2001). The development of new *Miscanthus* varieties that are regionally adapted as well having greater yield potential would greatly expand the economically viable growing region (Clifton-Brown & Lewandowski, 2000; Dong et al., 2019). The parental species of M. ×g, both MSA and MSI pose vast genetic resources to improve the production of more M. ×g varieties (Sacks et al., 2013). Therefore, exploring *Miscanthus* parental genotypes that are widely adapted to diverse environments will aid in breeding for improved M. ×g crosses.

Exploring the broader adaptation of *Miscanthus* genotypes requires their evaluation in multiple environments where they are assessed under diverse environmental conditions. This is important, especially for complex traits like biomass yield, where genotype-by-environment interaction (G×E) causes challenges by changing the ranking patterns of genotypes across different environments (Jarquin et al., 2021). Furthermore, breeders in the traditional *Miscanthus* breeding program require up to three years to collect trustable phenotyping data (Lewandowski et al., 2016). Considering the utmost goal of the breeding program to improve genetic gain, delayed phenotyping, and the extended evaluation period in METs can hinder early decision-making to release the most promising genotypes.

Genomic selection (GS) offers an alternative approach to utilizing whole-genome marker information to select superior genotypes at the early stage of the breeding program, thereby increasing genetic gain per unit of time (Meuwissen et al., 2001). It allows breeders to make faster decisions regarding selection without waiting for the phenotypic information of the individuals (Eggen, 2012). Briefly, GS initially requires two types of datasets: training (TRN) set and testing (TST) set, where the TRN set contains both phenotypic and genotypic data, and the TST set contains only the genotypic data (Isidro et al., 2015). Before implementing GS in real-life applications, it is necessary to evaluate the suitability of the data at hand by simulating prediction scenarios mimicking these realistic prediction problems of interest to breeders through different cross-validations (Haile *et al*., 2021).

These simulation studies require splitting the already observed data into TRN and TST sets under different cross-validation schemes, then calibrating the models using the TRN set and evaluating their predictive ability (PA) in the TST set (Cheng et al., 2021; McGaugh et al., 2021). Such that GS can predict the performance of the genotypes that were never observed in the given fields. Eventually, the results of the GP model are used to calculate the genomic estimated breeding values (GEBVs) of the lines based on their marker information (Hayes et al., 2009). Therefore, genotypes can be selected earlier in the breeding program, speeding the breeding cycle and saving the cost of evaluating them in fields (Krishnappa et al., 2021).

Dealing with multi-environment data, the efficiency of GP models is evaluated based on the trial basis PA by measuring the correlation between the predicted and the observed values within the same environment. In recent years, GP models such as the genomic best linear unbiased prediction (GBLUP), a convenient reparameterization of the penalized ridge-regression best linear unbiased prediction (rr-BLUP), have been considered as the gold standard approach (VanRaden, 2008; VanRaden et al., 2009; Endelman, 2011). In addition, GS have been implemented using the Bayesian paradigm, allowing different prior distributions for the marker effects and using Markov-Chain Monte Carlo (MCMC) for estimating the parameters (Wang et al., 2018).

While GBLUP/rrBLUP account for a common variance for all markers, under Bayesian approach, different variances can be modelled (Meuwissen et al., 2001, de los Campos et al., 2013). Moreover, GP models have been expanded to include G×E interaction to better identify high-performing genotypes across different environments. The STME models can integrate environment covariates (ECs) into the model to capture the interaction between these and genomic markers (Jarquin et al., 2014). Additionally, Lopez-Cruz et al., (2015) proposed a multi-environment GP model that accounts for G×E interaction by considering all the different contrasts between genomic markers and environments. While GS has been successfully implemented in different breeding programs using the above-mentioned GP models, these are limited to predicting only one trait at a time.

Alternatively, multi-trait GP models leveraging the genetic correlation between pairs of traits for improved PA have shown to outperform uni-trait GP models (Jia and Jannink, 2012) in some specific situations. While single-trait models analyze each trait separately, multi-trait GP models use shared genetic signals across traits. This approach provides advantages for traits with low heritability (Gill et al., 2021). Additionally, when breeders choose to focus on a single trait, they can also incorporate other secondary traits into the models to improve the overall PA for the primary traits (Velazco et al., 2019). Several studies have implemented GP-based multi-trait approach for breeding crops such as wheat, sorghum, soybean (Schulthess et al., 2018; Fernandes et al., 2018; Velazco et al., 2019; Persa et al., 2020).

Moreover, Osterman et al., (2024) found that the multi-trait model achieved better PA than single-trait models in red clover, particularly when the genetic correlation between traits was 0.5 or higher. As mentioned, increasing PA holds the potential to accelerate genetic gain for the superior genotypes hence the evaluation of the multi-trait GP models is of interest. Considering multi-trait GP models for single environment prediction requires observation of at least one of the traits for a genotype to produce significant improvements in PA (Persa et al., 2020).

In recent years, multi-trait models have been extended to the multi-trait multi-environment model (MTME) that allows breeders to capture trait correlations as well as G×E interaction (Montesinos-López et al., 2019a). The MTME approach can improve the PA of the GP models in different breeding programs (Gill et al., 2021). It is also advantageous for traits that are hard to observe in one environment, while correlated traits in other environments provide additional details. Recently, Montesinos-López et al. (2019b) proposed a Bayesian MTME (BMTME) model that accounts for the estimation of variance-covariance structure among genotypes, traits, and environments. This approach can predict multiple traits tested in different environments. Ibba et al. (2020) performed a simulated study using the BMTME model that outperformed single-trait models. Similarly, Guo et al. (2020) found an improvement in PA utilizing BMTME models over single trait models for agronomic traits in wheat.

The BMTME approaches implemented to fit the traditional multi-trait multi-environment model organize the phenotypic data into a matrix format where rows represent the genotypes and the columns the traits in particular environments, creating a two-dimensional matrix as an input while fitting the model. This format does not allow model fitting for specific situations when fewer or null combinations of genotypes across traits and environments are observed impeding the computation of covariance structures. For example, those cases where new genotypes are being predicted in unobserved environments (CV00).

In this study, all the observations across genotypes, traits, and environments are organized into a single vector. The entries of the rows correspond to a genotype in trait and environment combinations. The phenotypic data is organized into a single vector as opposed to the traditional MTME approach, where data for multiple traits is typically arranged into separate columns, while the genotypes are in the rows to create a matrix arrangement. While the implemented strategy contrasts with the conventional MTME approach, it allows leveraging information from related traits through variance-covariance structure derived from the incidence matrices. This approach enables the flexible integration of incomplete data, allowing for the inclusion of genotypes that lack information for certain traits across environments. Also, this can be particularly useful in GS when phenotypic information for the genotypes in a new environment is unavailable.

Although different GS approaches have been implemented in *Miscanthus* breeding programs to predict the performance of complex traits (Clark et al., 2019; Olatoye et al., 2019; Olatoye et al., 2020), no previous studies have focused on the use of GP-based MTME models for improved PA. *Miscanthus* is usually measured for biomass yield and several yield component traits (Clark et al., 2014; Clark et al., 2019). Applying the MTME approach in the *Miscanthus* breeding program for utilizing information across traits and environments could enhance the PA of complex traits. In this study, using both STME and MTME GP models we focused on predicting the performance of the genotypes in three situations: 1) predicting untested genotypes that are never observed for any trait across multiple environments 2) predicting genotypes for missing traits at a given environment and 3) predicting partially observed genotypes for at least one trait in some environments. The aims of the study were to *1*) implement both STME and MTME GP models using 336 *M. sacchariflorus* genotypes tested in three environments and scored for four traits, and *2*) contrast the performance between the MTME and STME models for predicting complex traits across multiple traits and environments in a *Miscanthus* population.

## Materials and methods

### Phenotypic data

The data set used in this study was obtained from Njuguna et al. (2023) where initially the phenotypic and genomic data were analyzed for 590 MSA genotypes that originated from East Asia. The trials were conducted in four environments [Sapporo, Japan by Hokkaido University (HU); Urbana, Illinois by the University of Illinois (UI); Chuncheon, South Korea by Kangwon National University (KNU) and Zhuji, China by Zhejiang University (ZJU)] during 2015, having between one to four replicates in each location in a randomized complete block (RCB) experimental design. The phenotypic data was collected in 2016 (year 2) and 2017 (year 3) for above-ground dry biomass and 15 yield component traits.

Best linear unbiased estimates (BLUEs) for each genotype-trait-location combination across years were provided. These BLUEs were used as an input for the analysis in this study. To achieve a complete dataset with no missing entries for any genotype, trait or environment, data was restricted to 336 genotypes at HU, KNU and ZJU. The four traits under analysis were dry biomass (YDY), culm node number (CNN), total culms (TCM), and average internode length (AIL). Thus, for the analysis, a total of 336 × 3 = 1,008 phenotypes per trait were used in this study.

### Genotypic data

The population was genotyped using restriction site-associated DNA sequencing (RAD-seq). The details of the library preparation are discussed by Clark et al., (2014). Briefly, the libraries were sequenced with Illumina HiSeq 2500 and 4000. The TASSEL-GBS pipeline was used for read alignment and SNP calling using the MSI reference genome (Bradbury et al., 2007; Mitros et al., 2020). Initially, the SNP dataset was filtered to include only SNPs with a minimum call rate of 70% and a minor allele frequency (MAF) of at least 0.05, resulting in a total of 268,109 SNPs. Subsequent filtering removed SNPs with more than 50% missing values and an MAF cutoff of less than 0.03. After applying this quality control, a total of 136,814 SNPs remained for analysis.

### Genomic prediction models

In this study, the performance of two single-trait multi-environment (STME) and three multi-trait multi-environment (MTME) GP models were evaluated for predicting four traits in three locations. The models were compared based on the prediction accuracies (PA) and mean square error (MSE) in different cross-validation scenarios. The details regarding the models and cross-validations are provided below:

### STME GP models with and without interaction

#### M1: E+L+G

Under the STME framework, the baseline model (**M1**) incorporates the main effects of the line (L), environment (E), and genomic factor (G). Here, G contains the information of the genomic markers in a form of a covariance structure. The main effects follow an independent and identical normal distribution. This base model can leverage information from shared genetic relationships learned from the training datasets to predict untested genotypes in the testing set. However, since the genotype-by-environment interaction (GEI) term is not included, it cannot predict changes in the rank of performance of genotypes across environments.

Consider that for a given trait 𝛾 (𝛾 = 1,2, …, 𝑡), 𝑦_𝑖𝑗_ represents the adjusted phenotypic observation of the 𝑖^𝑡ℎ^ (𝑖 = 1,2, …, 𝑛) genotype in the 𝑗^𝑡ℎ^ (𝑗 = 1,2, …, 𝐽) environment and is described as the sum of the common mean 𝜇, the random effect of the 𝑗^𝑡ℎ^ environment (𝐸_𝑗_), the random effect of the 𝑖^𝑡ℎ^ genotype (𝐿_𝑖_) as well as its corresponding random genomic effect based on marker SNPs (g_𝑖_), plus a random error term (𝜀_𝑖𝑗_) accounting for the non-explained variability. Here, g_𝑖_ is the linear combination between *p* markers (𝑋_𝑖𝑚_) and their corresponding marker effects (𝑏_m_) {*m* = 1,2,…,*p*} such that 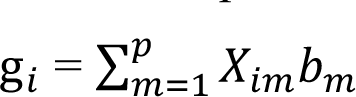, where 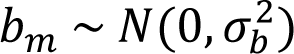 with 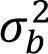 representing the respective variance component. Here, all the genomic information is compiled into a single vector 𝐠 = {g_i_} and by properties of the multivariate normal distribution 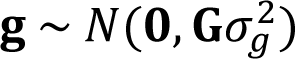, where 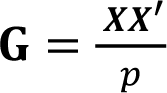 is the genomic relationship (VanRaden, 2008) whose entries describe similarities between pairs of individuals, 𝑿 is the standardized (by columns) matrix that of marker SNPs, and 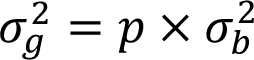 is the corresponding covariance component.

In addition, the model, the random terms 𝐸_𝑗_, 𝐿_𝑖_, and 𝜀_𝑖𝑗_ are assumed to be independent and identically distributed following normal distributions with mean 0 and a common variance for each component. Therefore, the model can be written as follows

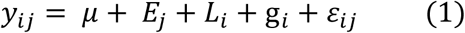

where,

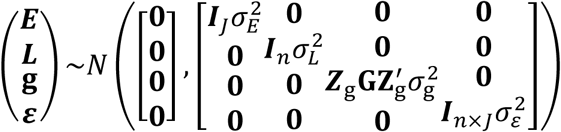

Here 𝒁_E_, and 𝒁_g_ are the design matrices connecting observations with environments and lines, and 𝑰 is the identity matrix of corresponding order the subindex, 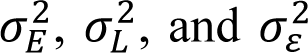 are the variance components associated with environments, genotypes, and the error term, respectively. This model enables the sharing of information between observed and unobserved individuals through the genomic relationship matrix **G** which helps to predict the performance of new genotypes.

#### M2: E+L+G+(G×E)

**M2** extends the base model **M1** by adding the interaction term G×E. This model enables different rankings of the genotypes across environments. Here the random effect term g𝐸_𝑖𝑗_ is added, referring to the interaction between the 𝑖^𝑡ℎ^ genotype and 𝑗^𝑡ℎ^environment. Here, all the interaction effects are compiled into a single vector such that g*E* = 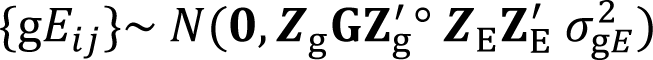 where 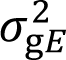 is the respective variance component, and “°” is the Hadamard product expressing the cell-by-cell multiplication of two matrices with the same dimensions (Jarquin et al., 2014). Thus, the model can be written as follows

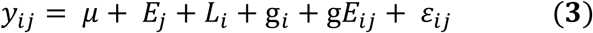

where,

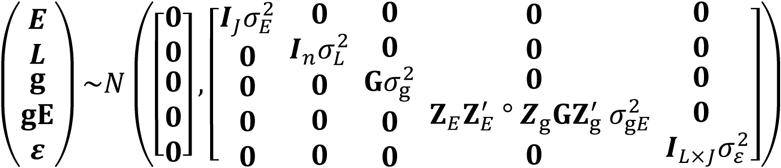

### MTME GP models with interactions

#### M3: E+T+(G×E×T)

**M3** contains the main effects of the environment (E), trait (T), and the complex three-way interaction between marker (G), trait (T), and environment (E). In the MTME models, traits can be correlated with each other, and it can result from both genetic and environmental factors. The G×E×T interaction can simulate trait-specific environmental influences. Here, the G is constructed the same way as described for the other models. However, this model does not explicitly account for the main effect of the markers. Here, the marker effect is only considered through its interaction with the environment and trait. The limitation of this model is that when selecting individuals, it is challenging to determine how much variance is attributed to genetic factors, and therefore, the genetic variance remains confounded with the environmental variance and trait variance.

Consider that 𝐲 is a vector with a length of 𝑛 × 𝐽 × 𝑡 (𝑛=number of genotypes, 𝐽=number of environments, 𝑡=number of traits). That is for 336 genotypes across three environments and four traits, 𝐲 vector contains a total observation of 4,032 (336 × 3 × 4). Hence, 𝑦_𝑖𝑗𝛾_ presents the phenotypic observation of the 𝑖^𝑡ℎ^ (𝑖 = 1,2, …, 𝐿) genotype in the 𝑗^𝑡ℎ^ (𝑗 = 1,2, …, 𝐽) environment for the 𝑡^𝑡ℎ^ (𝛾 = 1,2, …, 𝑡) trait. It can be explained as the sum of the common mean 𝜇, the random effect of the 𝑗^𝑡ℎ^environment (𝐸_𝑗_), the random effect of the 𝛾^𝑡ℎ^ trait (𝑇_𝛾_), the random interaction term among the genetic effect of the 𝑖^𝑡ℎ^ genotype (g_𝑖_) the 𝑗^𝑡ℎ^ environment and the 𝛾^𝑡ℎ^ trait, plus a random error term (𝜀_𝑖𝑗𝛾_). Like 𝐸_𝑗_, g_𝑖_, and 𝜀_𝑖𝑗_ in the previous models, 𝑇_𝛾_ is assumed to be independent and identically distributed. Briefly, in this model, 𝑇_𝛾_ presents a random vector of trait-specific effects across three environments. Unlike an unstructured variance-covariance matrix for the traits (usually used for the multi-trait models), we assume a homogeneous diagonal variance structure for each trait. Hence, each trait shares the common variance component likewise in 𝐸_𝑗_, g_𝑖_, and 𝜀_𝑖𝑗𝛾_.

Here, stacking the three way interaction effects into a single vector results in **g*E*T** = 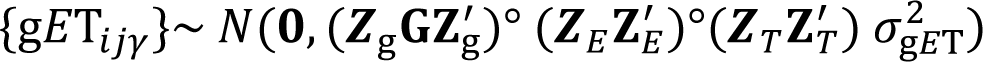 where 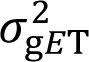 is the associated variance component. The explanation regarding the construction of **G** is described under model **M1**. Thus, the model can be written as follows.

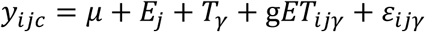

where,

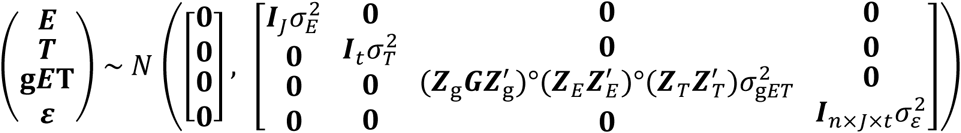

Here, 𝒁_𝑇_ is the incidence matrix connecting observations with the traits. All the other design matrices and error terms have been described before.

#### M4: E+T+G+(G×E×T)

**M4** is an extension of **M3** with the addition of the main effect of 𝑮. By modelling 𝑮 separately as a main effect, the model can obtain variance components due to genetic effects separately from the interaction. This model allows a better partition of the explained variance due to genetic differences only. The extension of model **M3** is conducted with the inclusion of the genetic effect of the genotypes (g_𝑖_), where 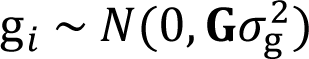 with 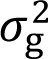 being the associated genetic variance. The construction of **G** is described under model **M1** and the rest of the terms are the same as in model **M3**. Thus, the model can be written as follows.

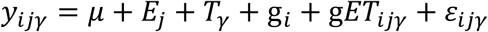

where,

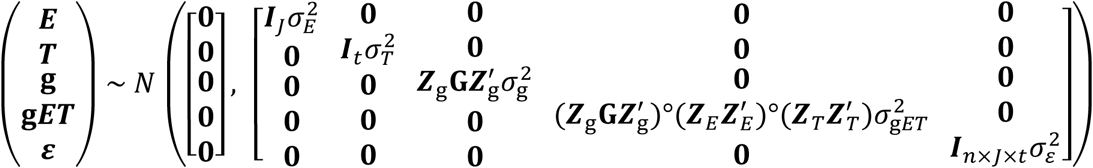

All the design matrices (𝒁_𝐸_, 𝒁_𝑇_ 𝑎𝑛𝑑 𝒁_g_) are consistent with those described in earlier models.

#### M5: E+T+G+(E×T)+(G×E)+(G×T)+(G×E×T)

This is the most elaborate model that accounts for all the interactions between environments and traits (𝐸 × 𝑇), genomic marker and environment (𝐺 × 𝐸), genomic marker and trait (𝐺 × 𝑇) and the three-way interactions among genomic marker, environment, and traits (𝐺 × 𝐸 × 𝑇) as well including all the main effects from model **M4** (E, T, and G). Each of the main effects has been described previously under different models. The interaction terms have been constructed by multiplying the variance-covariance structures of the corresponding main effects using the Hadamard product.

The inclusion of environment trait interaction (E×T) captures how trait performance varies across environments, G×E interaction accounts for the specific effect of genotypes in different environments, as earlier explained while G×T captures the genetic factor that affects multiple traits differently. Combining the previous results, the model can be written as follows

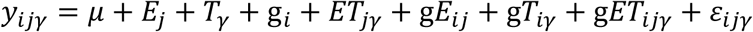

where,

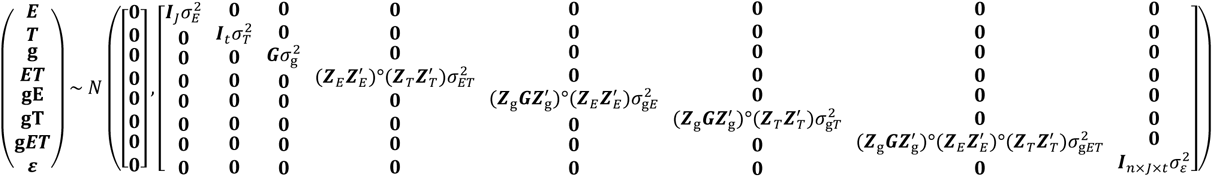

All the design matrices (𝒁_𝐸_, 𝒁_𝑇_ 𝑎𝑛𝑑 𝒁_g_) are as previously described. The symbol 𝜎^2^ denotes the associated variance components for each term. In this model, the inclusion of new interaction terms 𝐸 × 𝑇 and 𝐺 × 𝑇 influences the models to identify traits that are not sensitive to environmental changes and to select certain genotypes that are better suited for specific traits respectively.

### Cross-validations

The PA of both the STME and MTME models was evaluated using three cross-validation strategies, CV1, CVP, and CV2. These cross-validations mimic scenarios encountered in plant breeding. For all schemes, the dataset was divided so that prediction models were then trained using 80% of the data, leaving 20% of the data as a validation set to assess the performance of the models.

In CV1 the target is to predict the performance of the untested lines for all traits across all environments (**Figure 1**). To obtain a CV1 scheme, genotypes (not phenotypes) are randomly assigned to different folds. In this scheme, extra care was taken to allow the genotypes across all traits and environments to be assigned to the same fold. Therefore, when four folds (80% of the observations) were used to train the models, no information regarding the phenotypic data for any of the traits of the genotypes was available in the training set. The process of splitting data into training and test sets was repeated 10 times. CV1 poses major challenges for prediction as the training set lacks phenotypic information for any trait of the corresponding genotypes in the testing sets.

**Figure 1.**
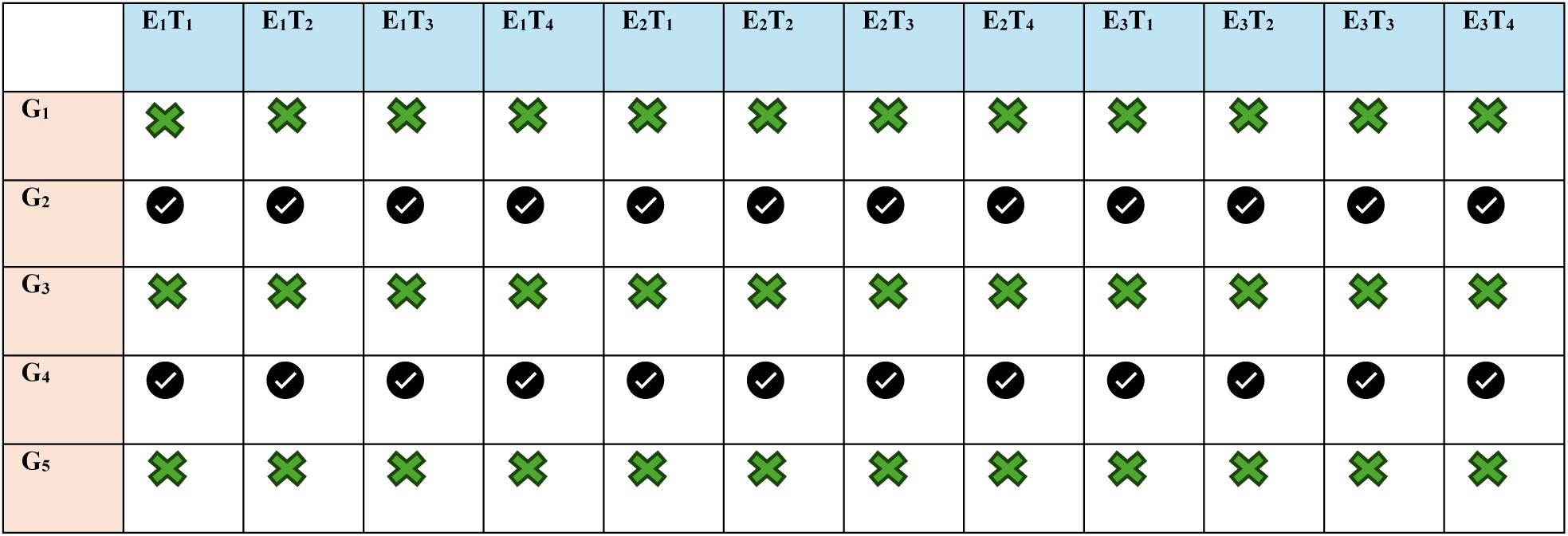
Schematic representation of CV1 for five exemplar genotypes (G_1_-G_5_) in three environments (E_1_, E_2_ and E_3_) across four traits (T_1_, T_2_, T_3_ and T_4_). The black symbol represents the genotypes having information for traits in environments. The green symbol shows the missing information of the genotypes for traits in environments.

CVP attempts to predict the performance of genotypes having missing trait information for a given environment (**Figure 2**) when information about the trait is available in other environments. For this, all traits from the same genotype in a specific environment were assigned to the same fold. For example, some genotypes for YDY, TCM, AIL, and CNN traits were assigned to fold 1 for the “CH” environment. When the other four folds (2-5) were used to test fold 1, the information for YDY, TCM, AIL, and CNN was unavailable for those genotypes in CH but was available in SP and ZJU.

**Figure 2.**
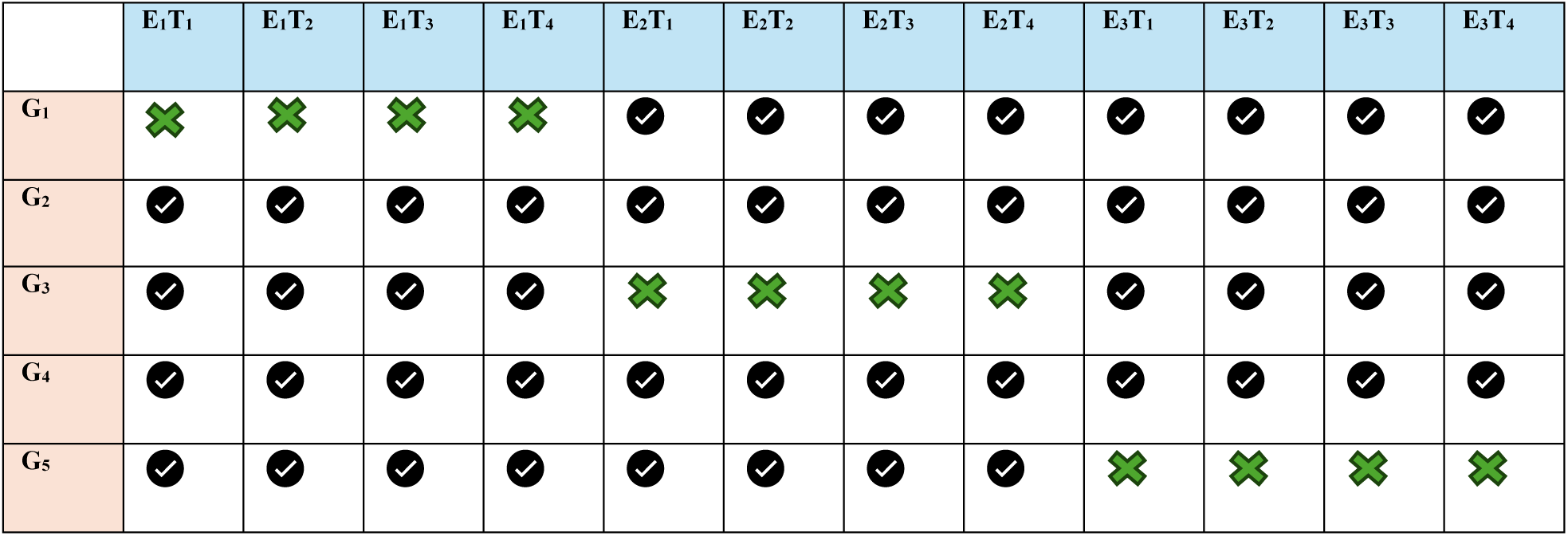
Schematic representation of CVP for five genotypes (G_1_-G_5_) in three environments (E_1_, E_2_ and E_3_) across four traits (T_1_, T_2_, T_3_ and T_4_). The black symbol represents genotypes that contain information for traits in various environments. The green symbol indicates the missing information for genotypes of traits in various environments.

Compared to CV1, CVP appears to be less complex for predicting genotype performance. This is because in CV1, the phenotypic information of individuals is completely masked across traits and environments, whereas in CVP, some partial information about the genotypes is still available across traits in other environments. In CVP, the models borrow information from those traits observed across other environments as well from traits observed for other genotypes in the same environment. In this case, partial data ensures a basis for prediction relative to the more daunting CV1 scenario.

The CV2 cross-validation scheme predicts the performance of the genotypes evaluated for at least one trait in some environments (**Figure 3**). To create a CV2 scheme, the observations across traits and environments are randomly assigned to five folds. Afterwards, four folds are used as the training set to calibrate the model, and the remaining fold serve as the testing set. Then the process repeated iteratively for each fold. This fivefold cross-validation technique was repeated ten times (replicates). The CV2 scheme is considered the easiest prediction scheme out of the three. Here phenotypic information from at least one trait for genotype is observed at each environment.

**Figure 3.**
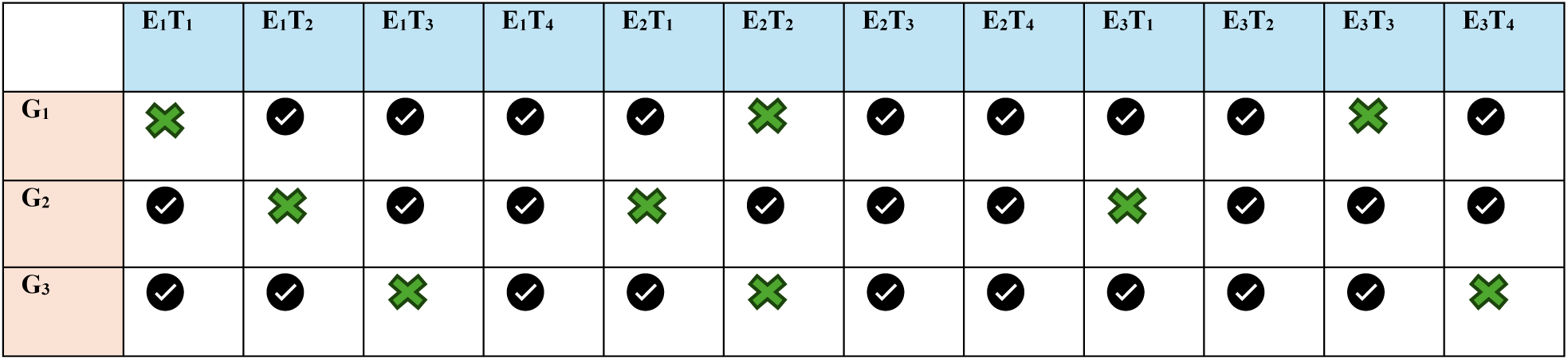
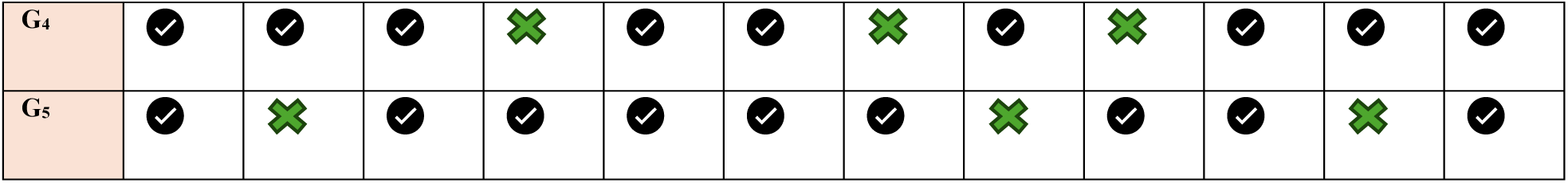
Schematic representation of CV2 for five genotypes (G_1_-G_5_) in three environments (E_1_, E_2_ and E_3_) across four traits (T_1_, T_2_, T_3_ and T_4_). The black symbol represents the genotypes having information for traits in environments. The green symbol shows the missing information of the genotypes for traits in environments.

For all cross-validations, the fold assignment was created in the MTME framework, resulting in a total of 4,032 observations across traits and environments. The same fold assignment was applied to the individual trait when implementing the STME models for the different cross-validation schemes. Under the STME framework, both CVP and CV2 are similar. For all schemes, the GP models are calibrated using both genomic and phenotypic data from the training set and these estimated effects are used to predict the testing set. Despite the framework (multi vs. uni trait) and the cross-validation schemes, the PA was measured as the Pearson correlation between predicted and observed values for each trait within the same environment.

## Results

The results are provided in three sections: (*a*) Prediction ability and mean squared error in the CV1 scheme; (b) Prediction ability and mean squared error in the CVP scheme; (c) Prediction ability and mean squared error in the CV2 scheme. The figures associated with each section are displayed in **Figures 4-6**, respectively.

**Figure 4.**
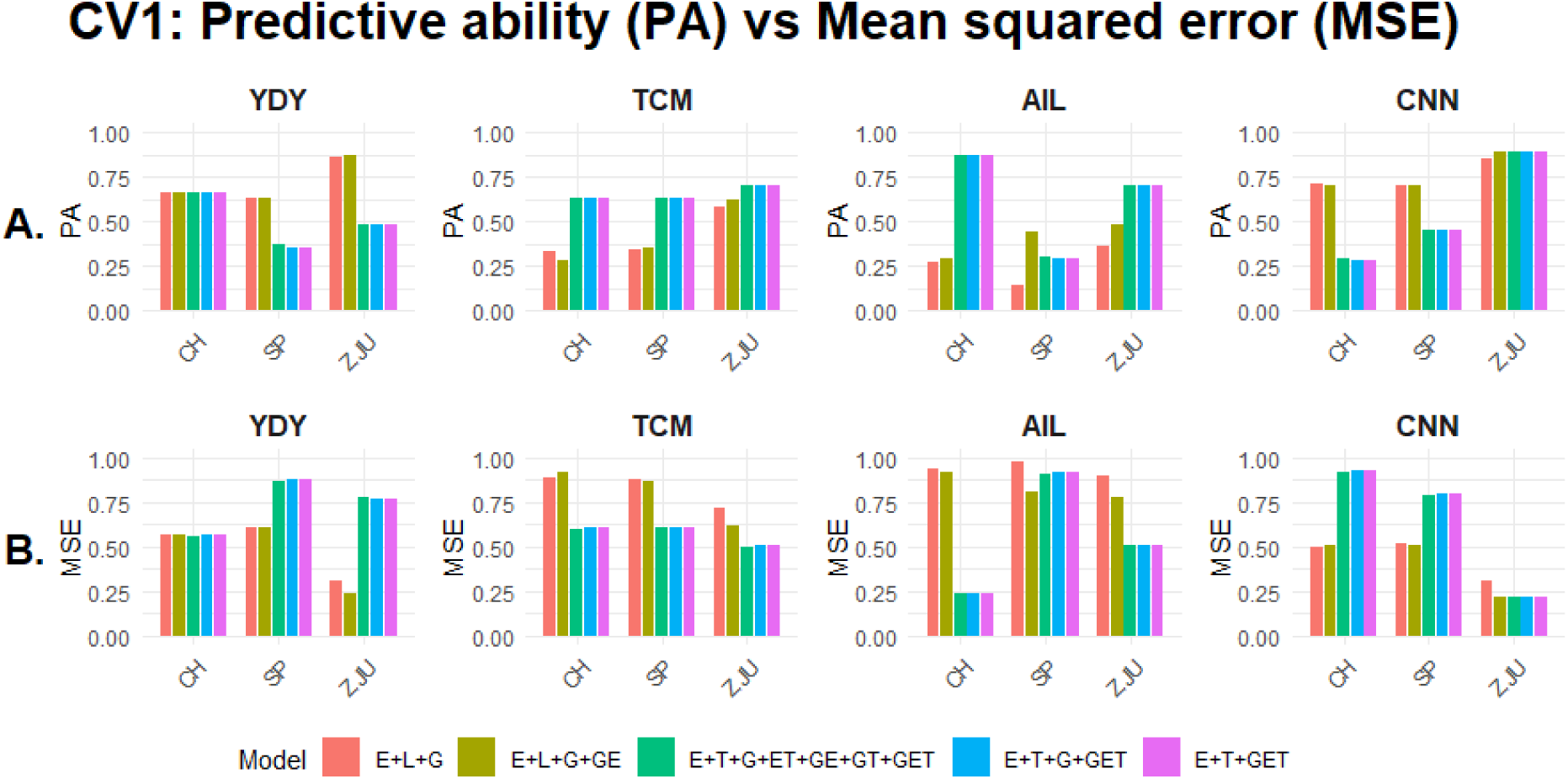
Average within-environment predictive ability (Panel A) and mean squared error (Panel B) in four *Miscanthus* traits (YDY: biomass yield, TCM: total culm number, AIL: average internode length, and CNN: culm node number) for two single trait multi-environment (E+L+G, E+L+G+GE) and three multi-trait multi-environment (E+T+G+ET+GE+GT+GET, E+T+G+GET, E+T+GET) GP models under the CV1 scheme across three environments (CH, SP, and ZJU). Different colors represent different GP models.

The PA and MSE in CV1, CVP, and CV2 were calculated separately for each environment (“CH”, “SP”, and “ZJU”) within each trait (YDY, TCM, AIL, and CNN). The predictions and MSE were compared under both single-trait multi-environment (STME) and multi-trait multi-environment (MTME) paradigms. The traits were scaled, which altered their original values; however, the relative MSE was used solely to compare model performance. Models with the highest predictions and lowest MSE values were considered the best performing.

### (a) Prediction ability and mean squared error in the CV1 scheme

**Figure 4** shows the results for the PA (**Panel A**) and MSE (**Panel B**) under the CV1 scheme across all traits and environments. For **YDY**, STME models performed better than MTME models, except in CH, where both models showed similar results with PA ∼ 0.69 and MSE ∼0.68. In SP, STME achieved a higher PA (∼0.67) and lower MSE (∼0.66), which was relatively 84% bigger than the PA and 16% smaller than the MSE obtained by the MTME models. At ZJU, STME again excelled MTME and achieved an increased PA (∼0.88 compared to ∼0.47) and smaller MSE, while MTME models showing 61-69% higher MSE.

In all three environments, MTME models consistently outperformed STME models for **TCM**. For example, in CH and SP, MTME models achieved a PA of ∼0.63, while STME models ranged between ∼0.28 and ∼0.36. At ZJU, MTME models had greater PA values of ∼0.70, compared with ∼0.58-∼0.62 in the STME models. Overall, MTME models had around 139% higher PA in CH and SP, and nearly 13% higher in ZJU, than STME models. Furthermore, MSE results also supported MTME, with lower values across all environments. In CH and SP, MTME models had an MSE of ∼0.65, whereas STME models had an MSE of ∼0.90. At ZJU, MTME models had an MSE of ∼0.50, in comparison to ∼0.67-∼0.73 under the STME frameworks.

Mixed patterns were observed for **AIL**, where MTME models were advantageous to STME models in CH and ZJU; however, no improvement was observed in SP. In CH, the MTME models had an overall PA of ∼0.88, outperforming the STME models by 238%. Similarly, MTME models achieved a constant MSE of ∼0.24, which was 220% lower than the MSE under the STME framework. At ZJU, the PA was higher in the MTME scheme, with an overall PA of ∼0.70. Comparatively, under the STME framework, the PA was ∼0.35 and 0.48 for M1 and M2, correspondingly. Hence, at ZJU, the PA of the MTME was nearly 100% and 46% greater than that of M1 and M2, respectively. Moreover, MTME models had a lower MSE of ∼0.50, which outperformed M2 model by 34% and the M1 model by 44%. Conversely, in SP, the STME M2 model was superior, with a larger PA (∼0.40) and smaller MSE (∼0.77), whereas the MTME had a 48% lower PA and a 15% higher MSE.

In addition, the worst performance was attained by STME M1 with a PA of ∼0.12 and an MSE of ∼0.99. Unlike TCM and AIL, where MTME models generally showed the best performance, STME results excelled MTME, particularly in CH and SP for the trait **CNN**. In CH, STME had a higher value, with a PA of ∼0.72 and an MSE of ∼0.50, whereas MTME showed a lower PA (∼0.26) and a higher MSE (∼0.77). In SP, STME again achieved a PA of ∼0.72 and a MSE of ∼0.50, which was more pronounced than MTME models, with a 44% decrease in PA and 52% increase in MSE. In ZJU, M2 model under the STME framework and all the MTME models performed similarly, with comparable PA (∼0.89) and MSE (∼0.22). In contrast, the M1 model had a lower prediction of ∼ 0.85 and a higher MSE of ∼0.33.

### (b) Prediction ability and mean squared error in the CVP scheme

The results for the CVP scheme across all four traits, with respect to PA and MSE, are shown in **Figure 5**. Across all traits, the trend was similar to that observed under CV1. The PA (**Panel A**) and MSE (**Panel B**) for **YDY** showed that the STME models consistently outpaced the MTME models in SP and ZJU, while both frameworks had similar PA in CH (∼0.68). In **SP**, STME achieved a PA of ∼0.67, a 25% improvement over MTME, and a 33% lower MSE. In **ZJU**, the M2 model under the STME paradigm achieved the highest PA (∼0.87), exceeding the STME M1 by 9% and the MTME models by 43%. In addition, MTME models achieved higher MSE, with values 200% and 114% higher than those of M1 and M2, respectively. In contrast, in **CH**, STME and MTME models performed similarly, with comparable PA (∼0.68) and similar MSE (∼0.56).

**Figure 5.**
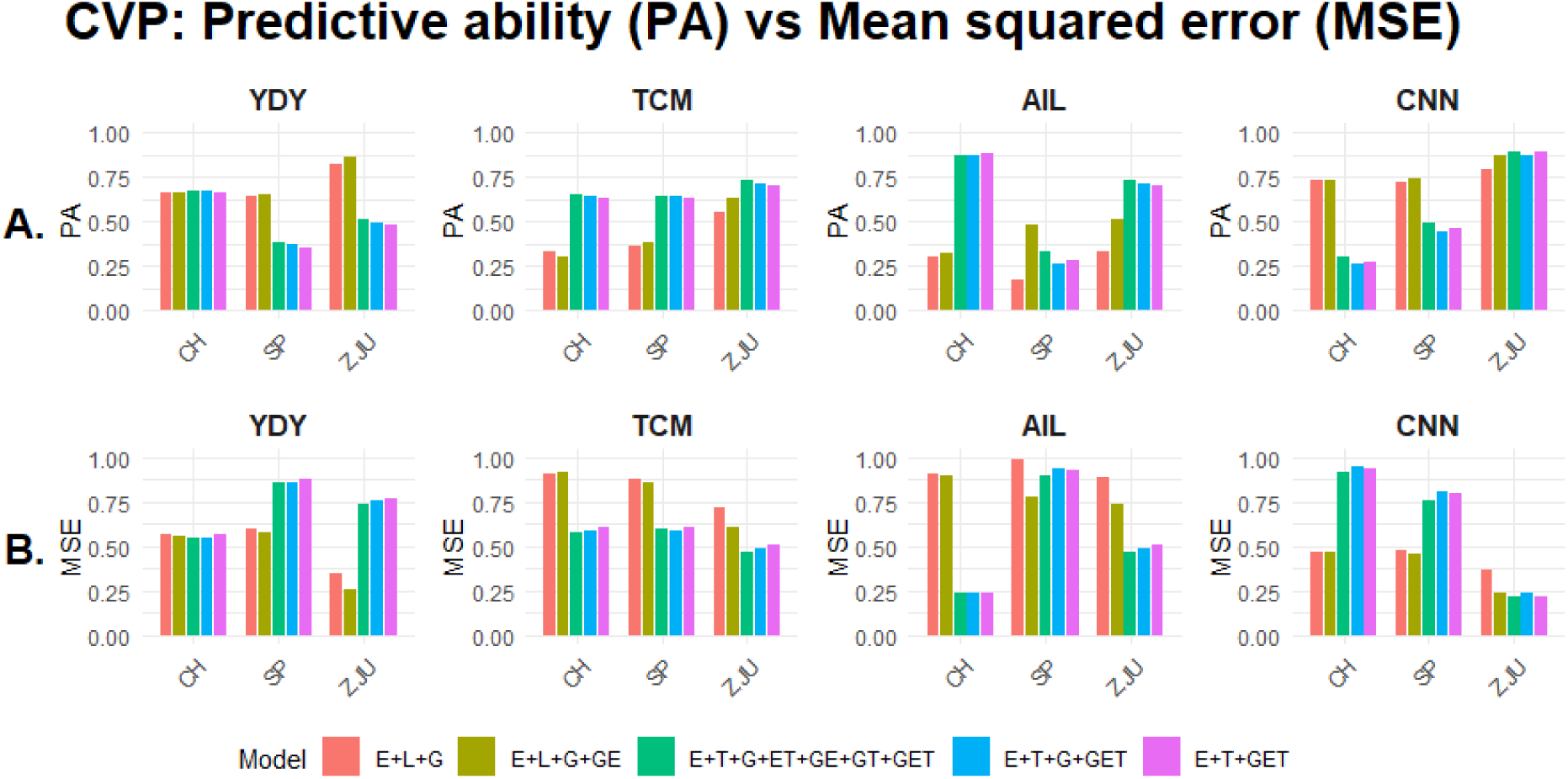
Average within-environment predictive ability (Panel A) and mean squared error (Panel B) in four *Miscanthus* traits (YDY: biomass yield, TCM: total culm number, AIL: average internode length, and CNN: culm node number) for two single trait multi-environment (E+L+G, E+L+G+GE) and three multi-trait multi-environment (E+T+G+ET+GE+GT+GET, E+T+G+GET, E+T+GET) GP models under the CVP scheme across three environments (CH, SP, and ZJU). Different colors represent different GP models.

Regarding **TCM**, MTME models consistently outpaced STME models across all environments, with maximum PA and minimum MSE. In CH, MTME models had a PA of ∼0.68 and a low MSE value of ∼0.65. Comparatively, STME models showed a decreased PA ranging from ∼0.27 to ∼0.32 and an increased MSE of ∼0.88. The results showed a 113% increase in PA and a 26% decrease in MSE for MTME compared to STME. A similar trend was present in SP, where MTME models had a PA of ∼0.68 and a MSE of ∼0.65, while STME models achieved lower PA (∼0.36-∼0.38) and higher MSE (∼0.88). This resulted in a 78% improvement in PA and a 26% reduction in MSE for MTME in SP. At ZJU, MTME models also had superior performance, obtaining PA, which was 8-35% higher than that of STME models. Similarly, MTME models achieved an MSE 24-30% lower than the STME models.

Considering **AIL**, the advantage of MTME models over STME models varied depending on the environment. In CH, MTME models exhibited strong performance, with a maximum PA of ∼0.88 and a minimum MSE of ∼0.23, whereas STME models performed the worst, achieving a PA that was 214% lower and an MSE that was nearly 275% higher than MTME. Similarly, at ZJU, MTME models had larger PA and smaller MSE, with PA values exceeding those of M1 and M2 in the STME paradigm by 37% and 10%, respectively. Additionally, the MSE values were 44% and 32% lower than those of STME. Conversely, in SP, the STME M2 model performed better, with a high PA of ∼0.48 and a low MSE. This PA was 220% higher than STME M1 and about 38% higher than the MTME models. Moreover, the MTME models had an MSE of ∼0.90, which was 10% lower than that of M1 and 20% higher than that of M2.

In contrast to TCM and AIL, STME models outperformed MTME models for **CNN** in both CH and SP. At ZJU, the performance of the MTME models was similar to M2 model under the STME framework. In both CH and SP, STME models attained similar PA values of ∼0.73, which was about 59% higher than the PA of MTME models in CH and 32% higher in SP. Similarly, STME models showed a decrease in MSE, being 88% lower than MTME in CH and 60% lower in SP. At ZJU, the STME M2 model and all MTME models had comparable PA of ∼0.88, which was 16% higher than that of STME M1. Likewise, the MSE values were similar, with the STME M2 and MTME models achieving nearly 38% lower values than STME M1.

### (c) Prediction ability and mean squared error in the CV2 scheme

The PA (**Panel A**) and MSE (**Panel B**) across all traits and environments under the CV2 are shown in **Figure 6**. Considering **YDY**, MTME models M4 and M5 had slightly better performance than the STME models in CH, with PA ranging from ∼0.68 to ∼0.70 and the MSE ranging between ∼0.50 and ∼0.54. The STME models and MTME M3 exhibited comparable performance, with a PA of ∼ 0.64 and an MSE of ∼0.60. In SP, the PA of STME models was 25-45% higher than that of MTME models. Considering MSE, STME models had an MSE of ∼0.64, whereas the MSE ranged from ∼ 0.72 to ∼ 0.88 for MTME models. At ZJU, the STME M2 model obtained the highest PA of ∼ 0.88 and the lowest MSE of ∼0.27, which outpaced STME M1 by 14% and the MTME models by 42%. In contrast, the MTME models attained the highest MSE values of ∼0.75, which were consistent across all models, showing a 200% increase in MSE compared to M2 and a 114% increase compared to M1.

**Figure 6.**
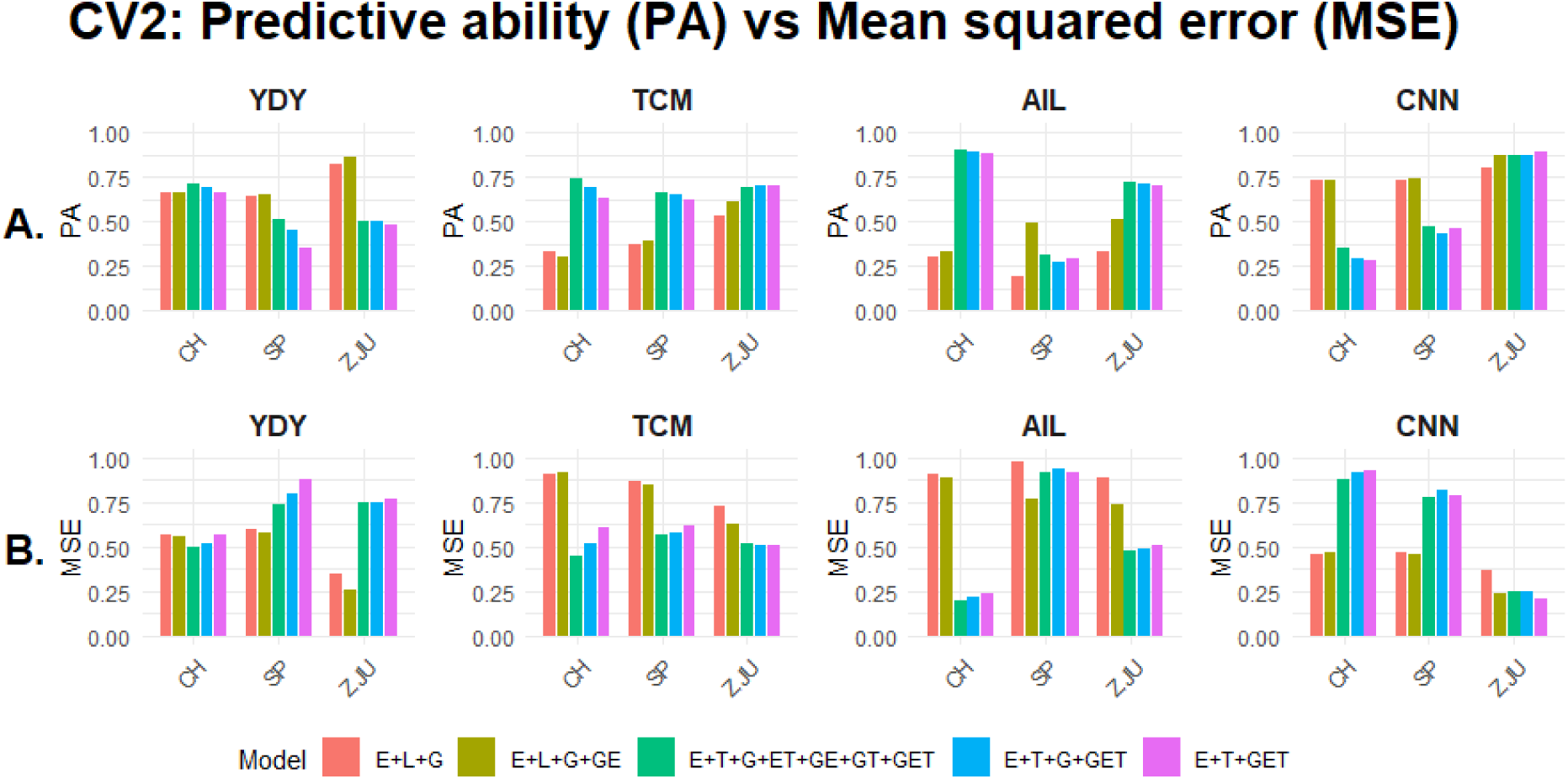
Average within-environment predictive ability (Panel A) and mean squared error (Panel B) in four *Miscanthus* traits (YDY: biomass yield, TCM: total culm number, AIL: average internode length, and CNN: culm node number) for two single trait multi-environment (E+L+G, E+L+G+GE) and three multi-trait multi-environment (E+T+G+ET+GE+GT+GET, E+T+G+GET, E+T+GET) GP models under the CV2 scheme across three environments (CH, SP, and ZJU). Different colors represent different GP models.

In **TCM**, the results aligned with the previous trend observed under the CV1 and CVP schemes, in which MTME models outperformed STME models in each environment. In CH, MTME models showed the highest PA values, ∼0.62 to ∼0.75, and the lowest MSE values ranging from ∼0.47 to ∼0.62. In comparison, both STME models had consistent minimum PA of ∼0.33 and maximum MSE value of ∼0.89. Similarly, in SP, MTME models obtained PA ∼0.66 and MSE between ∼0.58 and ∼0.62, whereas STME models had PA ∼0.39 and MSE ∼0.82. In ZJU, MTME models achieved a PA of ∼0.70 with an MSE of ∼0.50. The STME models performed the worst, with PA ranging between ∼0.54 and ∼0.63 and MSE between ∼0.65 and ∼0.70.

Likewise, in **AIL**, MTME models exceeded both STME models across all environments except SP. In CH, the MTME models achieved a PA of ∼0.88 and an MSE of ∼0.23, outperforming the STME models by 226% in PA and 74% in MSE. At ZJU, the MTME models obtained a PA of ∼0.71 and an MSE of ∼0.50, which exceeded the STME M1 and M2 models by 42-100% in terms of PA and by 31-46% considering MSE. Conversely, in SP, the M2 under STME was the best model, reaching a PA of ∼0.47, while the predictions under STME M1, MTME M3, and M4 ranged from ∼0.20 to ∼0.30, meaning that the M2 model performed around 56% better than the MTME models. M2 also had the lower MSE (∼0.76), while all other models (STME M1 and MTME models) had higher MSEs ranging between ∼0.90 and ∼0.95. Although MTME models performed slightly better than STME M1, the MSE values were still around 20% higher than STME M2.

Finally, for **CNN**, STME models showed superior performance to MTME models in CH and SP. In CH, STME models achieved a PA of ∼ 0.72 and an MSE of ∼0.43, presenting better performance than MTME models (PA ∼0.30 - ∼0.35 and MSE ∼0.90). Similarly, in SP, STME models had a higher PA (∼0.72) and a lower MSE (∼0.43), while MTME models achieved a PA between ∼0.42 and ∼0.44 and an MSE ranging from ∼0.75 to ∼0.77. At ZJU, both STME M2 and MTME models performed similarly, with PA of ∼0.88 and MSE of ∼ 0.25, while STME M1 showed slightly decreased PA (∼0.78) and an increased MSE (∼0.37).

## Discussion

One of the biggest challenges for using GP in breeding programs is to improve the PA of genotypic variation for complex traits that are highly influenced by environmental *stimuli*. This is because complex traits are prone to changing their response across environments due to G×E interactions. Therefore, it is not possible to select superior genotypes based solely on data from a single environment. Genomic selection has been successfully implemented in breeding programs to improve the PA of complex traits using multi-environmental data. In recent years, different GP models have been proposed to track G×E interactions by incorporating marker × environment interactions and have been shown to improve PA for the complex traits that are already observed in some environments (Jarquin et al., 2014; Lopez-Cruz et al., 2015; Crossa et al., 2017).

However, when the genotypes are not evaluated in any environment, it becomes challenging for the statistical models to leverage information of the other genotypes in different environments (Gill et al., 2021; Shahi et al., 2022). In addition, STME models that predict one trait at a time cannot use information from traits observed in other environments. Breeders evaluate genotypes for multiple traits for selection purposes, which provides opportunities to include them in the GP models (Shahi *et al*., 2022). Multi-trait (MT) GP models offer advantages over single-trait models by leveraging shared genetic information across traits (Sun et al., 2017; Shahi et al., 2022).

The extension of MT models to account for complex G×E interactions (MTME) adds benefits for predicting traits simultaneously across different environments (Montesinos-López et al. 2019a). The advantages of MTME models could be implemented in *Miscanthus* breeding programs since this crop is evaluated for multiple yield component traits across a diverse range of environments (Clark et al., 2014), enabling more comprehensive selection approaches to achieve enhanced genetic gain.

Several studies in *Miscanthus* breeding programs have utilized GP models to predict complex traits. Olatoye et al. (2019) used four uni-trait GP models and found the efficiency of GP models for predicting simulated traits with complex architecture. Clark et al., (2019) utilized uni-trait GP models in both single location and multi-location trials for predicting traits in *M.s sinensis*. Another study performed by Olatoye et al., (2020) used uni-trait GP model to predict the performance of a simulated progeny population by leveraging the genetic diversity from its parental population.

Moreover, the potential of GP models for predicting yield and yield component traits in *M. sacchariflorus* across multiple environments was demonstrated by Njuguna et al., (2023). However, despite *Miscanthus* being evaluated for many traits, no previous studies have explored the advantages of the multi-trait approach and contrast to the uni-trait GP models. To determine whether including multiple traits across environments improves the PA of the complex traits in the *Miscanthus* population, we evaluated the performance of two STME and three MTME GP models.

Our results suggest that the relative importance of STME and MTME models varied by traits and environments. MTME models consistently enhanced PA for TCM and AIL across most environments, showing successful implementation of MTME models. Conversely, for YDY and CNN, STME models often performed similarly or outperformed MTME models, indicating limited advantages of MTME modeling for these traits. Environmental factors also played a role, where MTME models exhibited more consistent performance in certain environments, whereas STME models (particularly M2) were advantageous in others. These findings suggest that the advantage of MTME models is both trait-specific and environment-dependent.

## Prediction scenario for CV1

In this study, it was found that in the CV1 scheme the MTME models performed better than STME models for TCM and AIL, but not for YDY and CNN. The MTME models with high PA also obtained lower MSE, highlighting better performance than the STME models. TCM and AIL had low phenotypic correlations between pairs of environments compared to YDY and CNN (**Figure S1**) whose showed similar correlation between pairs of environments. It implies that the performance of genotypes for these traits might vary across environments due to the influence of G×E. In that case, MTME models could borrow information by better capturing information across other genotypes observed in multiple environments, especially when low phenotypic correlations are present.

In the case of high phenotypic correlations, the models including interactions marginally improve the main effects models because the environments are alike. The improvement of MTME models in the CV1 scheme is noticeable as the genotypes are never observed in any environment. Previously, some studies reported no improvement of MTME models over STME models in the CV1 scheme (Jiang et al., 2015; Schulthess et al., 2016). As prediction in CV1 relies on other genotypes observed across environments, reduced PA in these studies could be associated with the absence of information from the environments where these genotypes are observed (Roorkiwal et al., 2018).

The improvements of MTME models over STME models were apparent for TCM in all environments and for AIL in two out of the three environments (CH and ZJU). MTME models had superior performance for traits exhibiting low phenotypic correlation across environments. However, in SP, low PA in MTME might be associated with AIL experiencing unique environmental conditions driven by plant-specific factors that cannot be measured (Hill and Mulder, 2010), and which MTME could not track.

Moreover, in SP for trait AIL, model M2 performed the best, while the lowest performance was observed for M1 compared to the MTME model. When the genotypes are never observed in CV1, it is challenging for **M1** model to pool information across environments. This is because M1 does not account for G×E interaction allowing models to capture specific signals in environments returning a weighted average value instead. On the other hand, in environments where plants exhibit variation due to non-recognizable micro-environmental factors (Raffo et al., 2023), it might be challenging for the MTME models to detect signals for those environments. Therefore, the reason why model M2 outperforms M1 and the MTME models could be due to its ability to capture the specific environmental variation for AIL in SP. The findings highlight the importance of environmental conditions on PA and the flexibility of M2 model in better handling within environment complex trait architectures.

For YDY, where using MTME models did not show advantages, STME models performed better in SP and ZJU, while in CH, all five models performed similarly. Moreover, both STME models with and without G×E interaction showed similar performance in all environments. However, the MSE value was the lowest for M2 in ZJU. In other words, there were no advantages of using G×E interaction term for predicting YDY in CH and SP, except in ZJU where the MSE value was low for M2. It is probably because YDY exhibited low G×E interaction due to high phenotypic correlation across different environmental pairs compared to TCM and AIL (**Figure S1**). While in ZJU, YDY showed variable response to environment-specific deviations, which the M2 model was able to capture. On the other hand, across CH and SP, the trait (YDY) remained stable, resulting in similar predictions for both STME models.

Considering CNN, at CH and SP, the STME models demonstrated similar and superior performance compared to the MTME models. Similar to YDY, CNN had high phenotypic correlation among all environments, unlike TCM and AIL (**Figure S1**). However, at ZJU, the performance of M2 was similar to that of the MTME models. This supports the fact that under high phenotypic correlations among environments, the G×E interaction term from M2 loses relevance compared to the main effects model. This was very clear for both YDY and CNN when comparing M1 vs. M2.

## Prediction scenario for CVP

In CVP, the performance of the models was similar to what was observed under CV1. The MTME models outperformed STME models for TCM and AIL but not for YDY and CNN. The similar performance of CV1 and CVP suggests that when a genotype is observed for all traits in other environments but is untested in a specific environment, the MTME models can still exploit patterns of the traits across multiple environments. In other words, for missing traits in one environment, observed trends of the genotypes across different environments benefit MTME models for predictions. Similarly to before, M2 outperformed all models in ZJU for YDY and in SP for AIL, indicating that this model well captured the impact of traits specific within-environmental variation in CVP.

## Prediction scenario for CV2

Apart from a slight improvement observed for YDY in CH using MTME models compared to STME models, the prediction scenario in CV2 presented similar patterns to those observed under the CV1 and CVP schemes. Previous studies have noted improvements in the CV2 scheme using MTME models (Wang et al., 2017; Lado et al., 2018; Bhatta et al., 2020; Gill et al., 2021). Because genotypes are observed for at least one trait in some environments but not in others, prediction under CV2 is less challenging. This cross-validation is useful for cases where some information of the genotypes is observed in the environment of interest. It might be the case, for example, that one trait is easier to measure than others. Some of these would involve destructive phenotyping techniques or high costs.

In these scenarios, partial information could be obtained for some traits but not others in the same environment for the same genotype. Or some traits could be strategically collected in some environments only. Unlike previous cross-validations, MTME slightly improved prediction for YDY in CH over STME models. This is possibly attributable to the proficiency of the models to integrate information regarding the overall performance of the genotypes across different environments.

## Conclusions

This study evaluated the potential of two single-trait multi-environment models and three multi-trait single-environment models for four yield component traits in *Miscanthus* population in three separate cross-validation strategies mimicking the real-life breeding program. An improvement of MTME models over STME models for trait TCM and AIL was observed in all cross-validation scenarios. This suggests that the MTME models can be implemented conveniently in the *Miscanthus* breeding program to improve PA for some traits while STME for others. In addition, additional advantages of MTME models in predicting YDY at a specific location under the CV2 scheme were identified. Further studies should consider different combinations of traits using the MTME models to improve PA in YDY.

It was also found that in some environments for particular traits, the M2 (E+L+G+G×E) model outperformed both the M1 (E+L+G) and the MTME models in CVP and CV2 schemes. It suggests that M2 is efficient in capturing the within-environment variation, while MTME benefits when environments have low phenotypic correlation among them. Overall, the PA obtained in this study using MTME approaches represents a new framework for utilizing GS in selecting complex traits within the *Miscanthus* population.

Finally, the approach presented here is an initial exploration of what can be done with MTME models for improving PA in *Miscanthus* population. The results presented here are quite satisfactory for improving PA in complex traits. Further research can include additional *Miscanthus* traits to enhance the prediction of yield component traits across all cross-validation scenarios. Although predictions for a completely new environment were not evaluated in the current study, this MTME approach provides flexibility for predicting unobserved genotypes in novel environments, which has been a challenge for similar implementations focused on estimating covariance structures. Future *Miscanthus* breeding programs should focus on the practical implementation of this MTME approach for improved genetic gain.

## Acknowledgments

This work was funded by the DOE Center for Advanced Bioenergy and Bioproducts Innovation (U.S. Department of Energy, Office of Science, Office of Biological and Environmental Research under Award Number DE-SC0018420). Any opinions, findings, and conclusions or recommendations expressed in this publication are those of the authors and do not necessarily reflect the views of the U.S. Department of Energy.

## Data availability

The prediction pipeline can be found in https://doi.org/10.6084/m9.figshare.31802728

## Appendix

**Figure S1.**
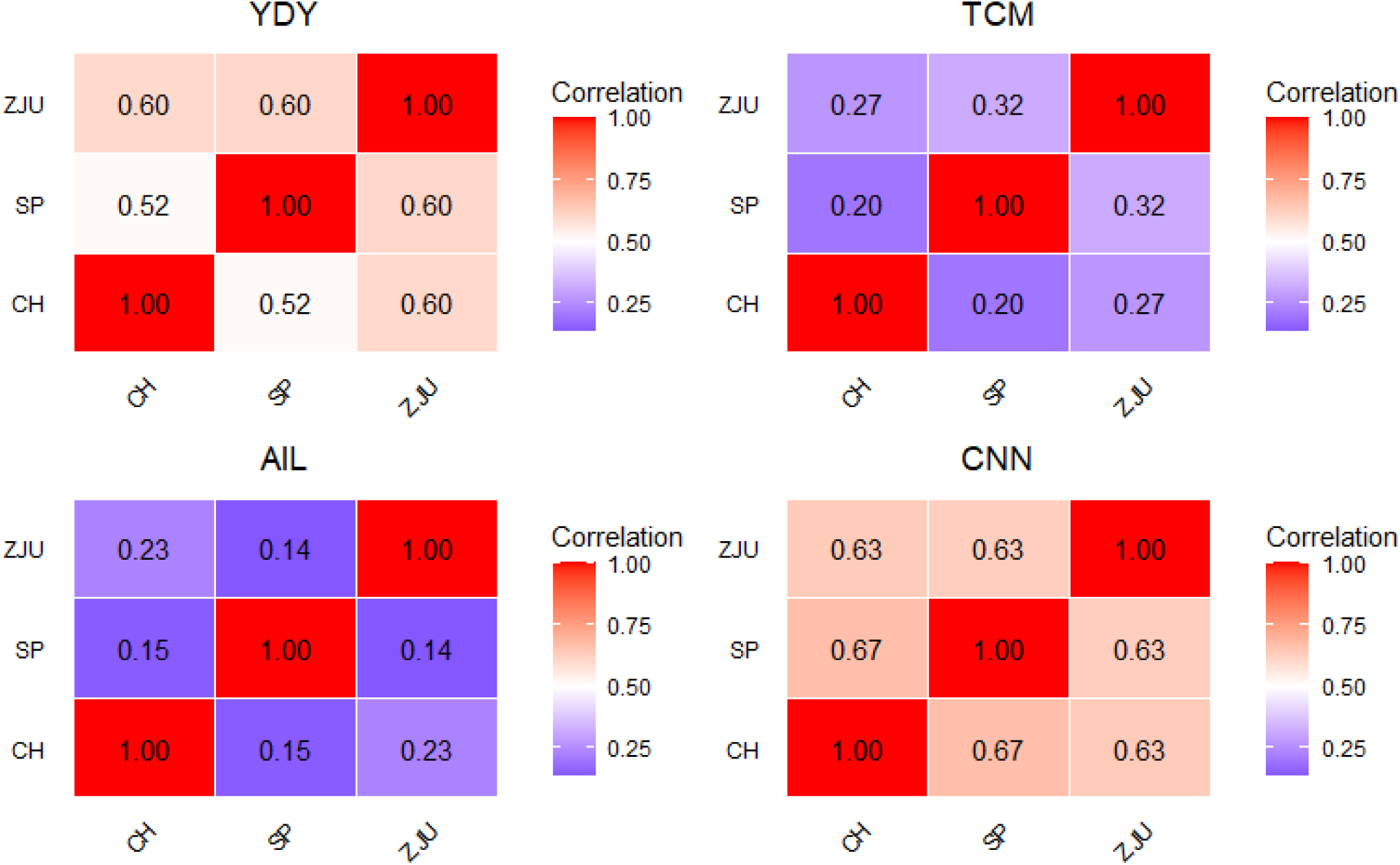
Phenotypic correlation between pairs of environments Sapporo, Japan by Hokkaido University (HU); Urbana, Illinois by the University of Illinois (UI); Chuncheon, South Korea by Kangwon National University (KNU) and Zhuji, China by Zhejiang University (ZJU) for a *Miscanthus sacchariflorus* population comprising 336 genotypes scored for biomass yield YDY, total culm number TCM, average internode length AIL, and culm node number CNN.

